# Targeted genomic surveillance of insecticide resistance in African malaria vectors

**DOI:** 10.1101/2025.02.14.637727

**Authors:** Sanjay C Nagi, Eric R Lucas, Faisal Ashraf, Trevor Mugoya, Edward Lukyamuzi, Shannan Summers, Calvin Yee, Christopher G Jacob, Harun Njoroge, Thomas Pemberton, John Essandoh, Martin Lukindu, Honorine Kaboré, Grégoire Sawadogo, Jessica Williams, Arjen E. Van‘t Hof, Anastasia Hernandez-Koutoucheva, Christina Hubbart, Kate Rowlands, Anna Jeffreys, Scott Goodwin, Naomi Park, Cristina Ariani, Alexander Egyir-Yawson, Sonia Goncalves, Shavanthi Rajatileka, Kirk Rockett, Victoria J. Simpson, Alistair Miles, David Weetman, Jonathan Kayondo, Tony Nolan, Martin J Donnelly

## Abstract

The emergence of insecticide resistance is threatening the efforts of malaria control programmes, which rely heavily on a limited arsenal of insecticidal tools, such as insecticide-treated bed nets. Importantly, genomic surveillance of malaria vectors can provide critical, policy-relevant insights into the presence and evolution of insecticide resistance, allowing us to maintain and extend the shelf life of these interventions. Yet the complex genetic architecture of resistance, combined with resource constraints in malaria-endemic settings, have thus far precluded the widespread use of genomics in routine surveillance. Meanwhile, stakeholders in sub-Saharan Africa are moving towards locally driven, decentralised generation of genomic data, underscoring the need for standardised and robust genomics workflows. To address this need, we demonstrate an approach to targeted genomic surveillance in *Anopheles gambiae s.l* with Illumina sequencing. We target 90 genomic loci in the *Anopheles gambiae s.l* genome, including 55 resistance-associated mutations and 35 ancestry informative markers. This protocol is coupled with advanced, automated software for accurate and reproducible variant analysis. We are able to elucidate population structure and ancestry in our cohorts and accurately identify most species in the *An. gambiae* species complex. We report frequencies of variants at insecticide-resistance loci and explore the continued evolution of the pyrethroid target site, the *Voltage-gated sodium channel.* Applying the platform to a recently established colony of field-caught resistant mosquitoes (Siaya, Kenya), we identified seven independent resistance-associated variants contributing to reduced efficacy of insecticide-treated nets in East Africa. Additionally, we leverage a machine learning algorithm (XGBoost) to demonstrate the possibility of predicting bioassay mortality using genotypes alone. This achieved high accuracy (75%), demonstrating the potential of targeted genomics to predictively monitor insecticide resistance. Together these tools provide a practical, scalable solution for resistance monitoring while advancing the goal of building local genomic surveillance capacity in sub-Saharan Africa.

## Introduction

Insecticidal interventions remain the cornerstone of malaria vector control, however, sustained control is challenged by the emergence and spread of insecticide resistance among the *Anopheline* vectors^1^. Addressing the problem of escalating resistance is vital to maintaining progress towards malaria elimination, which has stalled in recent years^2^. New active ingredients have recently become available, including long-lasting insecticidal nets (LLINs) with dual active-ingredients, such as pyrethroid-pyrrole (chlorfenapyr) and pyrethroid-synergist (PBO) combinations. After promising epidemiological results in randomised-controlled trials^3–6^, dual chlorfenapyr and PBO nets already make up the vast majority of nets delivered to sub-Saharan Africa^7^. Their widespread deployment raises concerns about replicating the earlier over-reliance on pyrethroid-only nets over the past two decades; monitoring wild vector populations for existing and newly emerging resistance to these novel active ingredients will be crucial to extend their longevity.

Genomic surveillance has emerged as a pivotal approach to understanding the spread and evolution of pathogens in many infectious disease systems (for example, *SARS-CoV-2*^8^, *mpox*^9^, *mtb*^10^) and offers great potential to help mitigate insecticide resistance in malaria vectors. By monitoring known and identifying new resistance-conferring mutations before they become widespread, control programs can implement proactive targeted interventions, such as rotating insecticides, deploying alternative active ingredients, or prioritising resistant vector populations for intensified monitoring. These measures help limit the spread of resistance and sustain the efficacy of existing vector control strategies. Therefore, a thorough understanding of the link between resistance genotype and phenotype is essential to devise mitigation strategies, however, current phenotype-based methods for the surveillance of insecticide resistance are both labour-intensive and imprecise^11,12^. Whole genome sequencing has revealed insights into mosquito biology^13^ and many genes and mutations in the *An. gambiae* genome involved in insecticide resistance^14–19^. Although whole genome sequencing is powerful, the relatively large size of mosquito genomes (*An. gambiae ≈* 278Mb) makes targeting specific genomic loci a cheaper and more tractable option for routine surveillance activities.

In this study, we develop a cost-effective approach to targeted genomic surveillance in *Anopheles gambiae sl*. We introduce Ag-vampIR: the *<u>A</u>n. <u>g</u>ambiae* <u>V</u>ector <u>A</u>mplicon <u>M</u>arker <u>P</u>anel for Insecticide <u>R</u>esistance, which contains 80 amplicons targeting 90 SNP markers, designed for Illumina sequencing platforms. The markers are selected to target known insecticide resistance loci and ancestry informative markers for the resolution of species within the *An. gambiae* complex. We also present open-source software, AmpSeeker, a computational workflow designed for the comprehensive analysis of any Illumina amplicon sequencing data. Leveraging the power of Snakemake^20^, AmpSeeker streamlines the entire analytical pipeline, from raw data processing to variant calling and downstream analyses, and generates a local webpage for users to conveniently explore the results. Our study presents the development and validation of this workflow alongside the *An. gambiae s.l.* Ag-vampIR amplicon sequencing panel. Together, these tools aim to enhance malaria vector surveillance through scalable, efficient, and reproducible monitoring of insecticide resistance.

## Results

### Platform and protocol development

To perform targeted genomic surveillance of *Anopheles gambiae sl,* we developed an Illumina amplicon sequencing panel, Ag-vampIR, and a library preparation protocol. The Ag-vampIR panel is based on the *An. gambiae* AgamP4 PEST reference genome and targets 90 SNP markers in total, split over 82 amplicons (Figure 1A). It incorporates major known insecticide resistance loci (55 SNPs in total), including target-site mutations, such as *Kdr, Ace1* or *Rdl*, markers for metabolic resistance, or SNPs that tag selective sweeps prevalent in the genomes of wild specimens from the *Anopheles* 1000 genomes project (Ag1000G) (Supplementary Data 1, Supplementary Text 2). Thirty-five genomic targets are ancestry informative markers to allow the accurate resolution of species, and we include two amplicons at the intron 4–exon 5 boundary of *doublesex*, the target of a recently developed population suppression gene drive ^21^. Each amplicon is approximately 200 bp long, and our method makes full use of both known and novel SNPs present on Ag-vampIR amplicons. The laboratory protocol utilises an indexing PCR reaction and a combination of 96 custom i7 adaptors and 16 i5 adaptors^22^, to give a theoretical maximum of 1536 samples which can be multiplexed on a single flow cell. The entire amplicon sequencing protocol is summarised in Figure 1B.

**Figure 1.**
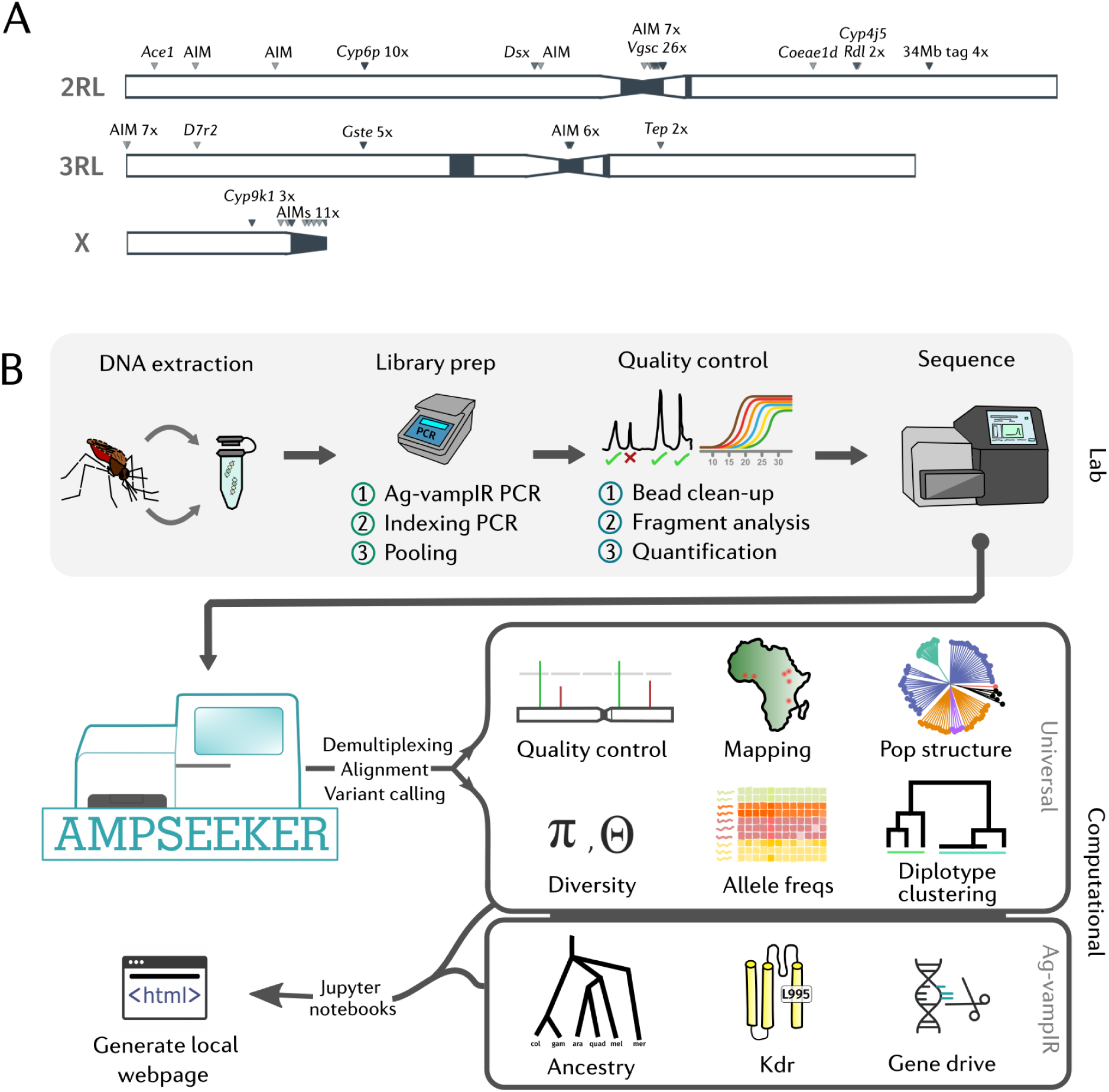
Ag-vampIR and AmpSeeker. **A)** A diagram of the genomic regions and SNP markers targeted by the Ag-vampIR panel (AgamP4 PEST reference genome). Heterochromatic regions are coloured black (for locus details see supplementary data 1). **B)** Schematic overview of the laboratory protocol and bioinformatic pipeline for targeted genomic surveillance. Upper: the laboratory and library preparation protocol. Lower: the AmpSeeker modules and computational workflow.

### Pipeline development

To enable reproducible, decentralised analysis of the amplicon data, we developed an open-source computational pipeline, AmpSeeker. The pipeline is written in Snakemake ^20^, and facilitates automated, end-to-end analysis. AmpSeeker can be applied to any Illumina amplicon sequencing data, with additional optional analyses for the Ag-vampIR panel. It is modular in architecture, allowing the analysis of further panels to be easily incorporated. Analyses are written in Python, the most popular programming language in the life sciences. The workflow is convenient to use and utilises Jupyter notebooks combined with Jupyter Book to generate a local web page of results for easy data exploration. Throughout this manuscript, we display outputs of the AmpSeeker pipeline.

### Sample collections and sequencing

We collected *An. gambiae s.l* mosquitoes from four cohorts across sub-Saharan Africa; two field populations; Obuasi, Ghana (2017), and the Upper River Region, The Gambia (2011), and from two laboratory strains, Siaya (Western Kenya, established in 2023) and VK7 (Burkina Faso, 2014) (Supplementary Table 1). These cohorts were selected to provide a diverse mixture of ancestries and resistance profiles with which to evaluate the targeted surveillance platform. We sequenced 542 samples in total on an Illumina MiSeq with a reagent kit V2.

### Quality control

Following sequencing, the MiSeq output was imported into AmpSeeker, which performs demultiplexing followed by alignment to the reference genome with bwa mem ^23^, and variant calling with bcftools mpileup ^24^. Following variant calling, we perform sample quality control; removing samples with low depth, and samples with extreme levels of heterozygosity or outliers in a principal components analysis (PCA). This resulted in a reduction from 542 samples to 467.

We then evaluated the accuracy of AmpSeeker’s variant calling, using whole-genome sequence data as a gold standard. 40 individuals (Obuasi, Ghana) were sequenced to 30x as part of the Ag1000g project, and previously analysed as part of a genome-wide association study ^14^. We examined the concordance between genotypes for these 40 samples at 11 variant read depth thresholds (Supplementary Figure 1). At 20X and above, we observe highly concordant variant calls (> 99.3%), demonstrating that our method is reliable and accurate.

### Taxon analysis and population structure

The *An. gambiae* species complex contains seven recognised species and at least four additional cryptic taxa, with varying contributions to malaria transmission^25–27^. Species within this complex are morphologically identical, underlying the importance of molecular methods to delineate the ancestry of an individual. Laboratory-based methods PCR assays^28,29^ are used to resolve species, however, these methods rely on single loci and cannot resolve cryptic taxa. Methods that rely on single loci also have limited power to detect hybridisation between species, a phenomenon which can occur in areas of sympatry. In order to accurately identify species, we developed an ensemble of three complementary methods: (1) an agnostic method based on ancestry informative markers (AIMs), (2) a PCA + SVM classifier and (3) an XGBoost classifier - both utilising reference data from 7790 published whole-genome sequences from the Ag1000g (see Methods). Taxons were assigned where at least two methods agreed, otherwise samples were designated as ‘unassigned’. Our AIM approach targets thirty-five AIMs from those published in Miles *et al*.^13^, allowing species delineation between *An. gambiae* and *An. coluzzii* at a greater resolution than traditional methods. AIMs are SNPs which show marked frequency or fixed differences between species and therefore give insights into the ancestry of a specimen. The PCA and XGBoost approaches allow us to expand our taxonomic analysis to other species in the *An. gambiae* complex whilst accounting for potential missing genotypes. To evaluate population structure alongside our taxonomic analysis, we built neighbour-joining trees and performed PCA.

Our analysis of population structure and ancestry revealed distinct patterns across sampling cohorts (Figure 2). The neighbour-joining tree and principal component analysis (Figure 2B, Supplementary Figure 2) separated our samples into five distinct clusters, which aligned with both geographic origin and assigned taxa. These clusters included: a large group of *An. arabiensis* from The Gambia, the Siaya colony samples (potentially the Pwani molecular form^27^), a cluster of *An. gambiae* primarily from Obuasi with four samples from The Gambia, and two clusters of *An. coluzzii* comprising the VK7 lab colony and a subset of the Gambian samples. Taxon assignments can be found in Supplementary Table 5 and Supplementary Data 3.

**Figure 2.**
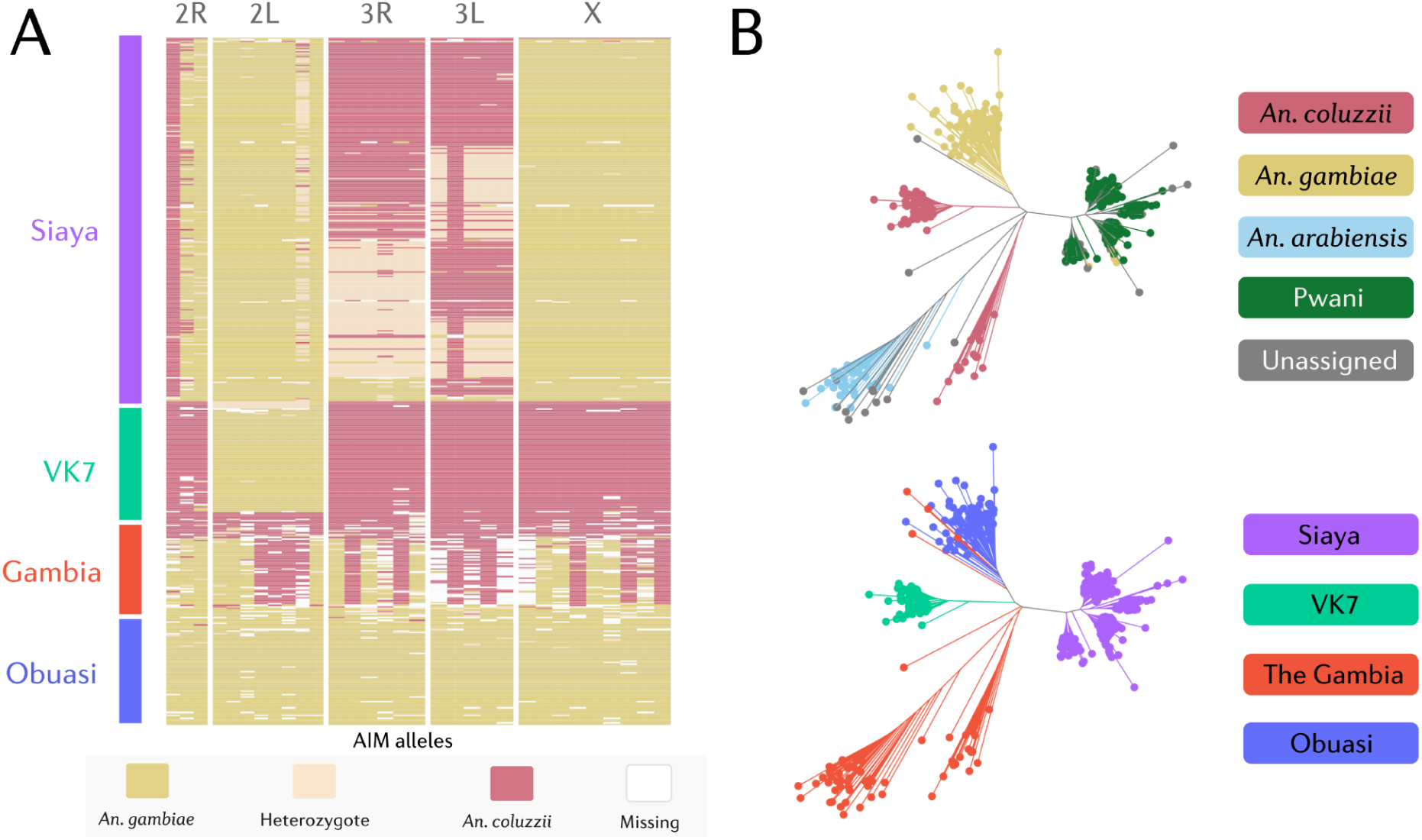
Ancestry and population structure. A) A heatmap of *An. gambiae* vs *An. coluzzii* ancestry informative marker (AIM) genotypes. Each row of the heatmap is a different sample, and each column a different AIM. AIMs are coloured according to the ancestral allele (gold: *An. gambiae*, maroon: *An. coluzzii*, and beige: heterozygote). B) A neighbour-joining tree of the 467 QC-passed amplicon sequencing samples coloured by sample taxon according to consensus taxon calls (upper) and sample location (lower).

The AIM analysis provided further insights into the ancestry of these samples (Figure 2A). The cohort from Obuasi, Ghana, contains *An. gambiae* s.s ancestry alone, in correspondence with taxon calls from the Ag1000g project. All Obuasi samples were correctly assigned as *An. gambiae* by our ensemble taxon caller. The VK7 strain of *An. coluzzii* appears to have AIMs consistent with *An. coluzzii* ancestry, apart from eight AIMs on the 2L chromosomal arm. The AIMs on this contig are clustered near the beginning of the arm, close to, or within the *Voltage-gated sodium channel (Vgsc)*, and the *An. gambiae*-like AIM patterns seen here are due to VK7 harbouring the 995F F1 *Kdr* haplotype (see *Vgsc* analysis below). This haplotype confers resistance to pyrethroid insecticides, and has introgressed from *An. gambiae* into *An. coluzzii* in West Africa^30^. The ensemble caller called all VK7 individuals as *An. coluzzii*.

The Gambia cohort contains mostly (*n*=45) a subgroup with an apparent mix of *An. gambiae* vs *An. coluzzii* ancestry, but also 17 samples with predominantly *An. coluzzii* genotypes and 4 with *An. gambiae* genotypes. While mixed patterns of *An. gambiae* vs *An. coluzzii* AIMs could indicate recent hybridisation or cryptic taxa (such as the recently reported Bissau molecular form^25^) both our XGBoost classifier and the principal components analysis predicted that the ‘mixed’ ancestry samples were in fact *An. arabiensis* (Supplementary Figure 3A). All three taxa are known to inhabit the Upper River Region of The Gambia ^31^.

The Siaya colony presented an intriguing pattern: ancestry consistent with *An. gambiae* on chromosome 2 and the X chromosome, but *An. coluzzii*-like ancestry on chromosome 3 (Figure 2A). This is surprising, as despite a recent finding of *An. coluzzii* in northern Kenya, this species has never been reported in Siaya county, western Kenya. Laboratory contamination is also unlikely, as the Siaya colony had never been maintained in the same insectary as a strain of *An. coluzzii.* Given the unusual AIM patterns in Siaya mosquitoes originating from Western Kenya, we hypothesised that this may relate to the origin of the AIMs themselves, which were ascertained from an earlier study of West African mosquitoes. To investigate this further, we evaluated the performance of these AIMs using Ag1000g data from East Africa specifically (Kenya, Uganda, Tanzania, 2880 samples) and found that most AIMs remained highly discriminative (>97% concordant) for species identification in the region. Further analysis with PCA (Supplementary Figure 3B) showed Siaya clustering in close proximity to both the *An. gambiae* cluster and adjacent to the cluster of the recently described *Pwani* molecular form (labelled *Gcx3*)^27^. In Ag1000g data, the *Pwani* form exhibits mixed AIM patterns on autosomes, in comparison to samples assigned to *An. gambiae*, which consistently show *An. gambiae-*like AIMs on the autosomes (Supplementary Figure 4). *Pwani* was thought to be limited to the coast of East Africa, however, a single individual female mosquito assigned as *Pwani* in the Ag1000g was collected in Muleba, Tanzania, on the West coast of Lake Victoria in 2015 (Supplementary Figure 5), suggesting that this subspecies may also exist inland.

### Surveillance of the Voltage-gated sodium channel

The Voltage-gated sodium channel (*Vgsc*) is the target of pyrethroids and DDT and across insect taxa has been the site of rapid evolution ^32^. In *An. gambiae*, phenylalanine and serine mutations at the 995 codon, known as *kdr*, lower the binding potential of the channel, enabling mosquitoes to survive exposure ^33,34^. Both 995F and 995S mutants have arisen independently on multiple occasions, with some origins remaining in more localised areas and others spreading over much of sub-Saharan Africa ^18^. We calculated allele frequencies at known SNPs within the *Vgsc* across our sample populations (Supplementary Figure 6). We also used tagging SNPs targeted by the Ag-vampIR panel to estimate the origins of *Kdr* haplotypes in our data (Supplementary Table 5).

In the Siaya strain from Western Kenya, we observed a mixture of 995F (65.9%) and 995S (33.7%) alleles, consistent with recent evidence of the westward spread of 995F in *An. gambiae* across central and East Africa^35^. Siaya contained multiple origins of the 995S allele (S1=31% and S3=1.5%) and a single origin of 995F, the F2 haplotype. VK7 were fixed for 995F (100%), with no evidence of the recently described V402L/I1527T haplotype, in agreement with its low frequency in West Africa during the time period of the colony’s establishment ^18,36^. The VK7 995F *Kdr* alleles all belonged to the major F1 haplotype. In The Gambia, *An. arabiensis* mosquitoes contain the 995S mutation at 16% frequency. We were not able to identify the origin of this mutant, which could suggest a novel origin in this sub-species. In samples assigned as *An. gambiae* from The Gambia, mosquitoes were fixed for 995F, with 83% of those samples belonging to the F1 haplotype, and the other 17% undetermined. In *An. coluzzii* from The Gambia, most samples were assigned as wild-type, with a small proportion (7.6%) of 995F alleles.

To visualise haplotypes and mutations at the *Vgsc* in more detail, we performed diplotype clustering at this locus, plotting a dendrogram alongside sample heterozygosity and any *Vgsc* mutations present in our data (Figure 3A). In agreement with the *Kdr* origins analysis, we identified two distinct haplotype backgrounds associated with the 995F mutation. The VK7 and Obuasi cohorts shared a common 995F haplotype (F1) characterized by multiple secondary mutations across the *Vgsc* gene region, including T791M, N1570Y, A1746S, A1853I, and P1874S/L. In comparison, Siaya contains the F2 haplotype lacking these additional variants.

**Figure 3.**
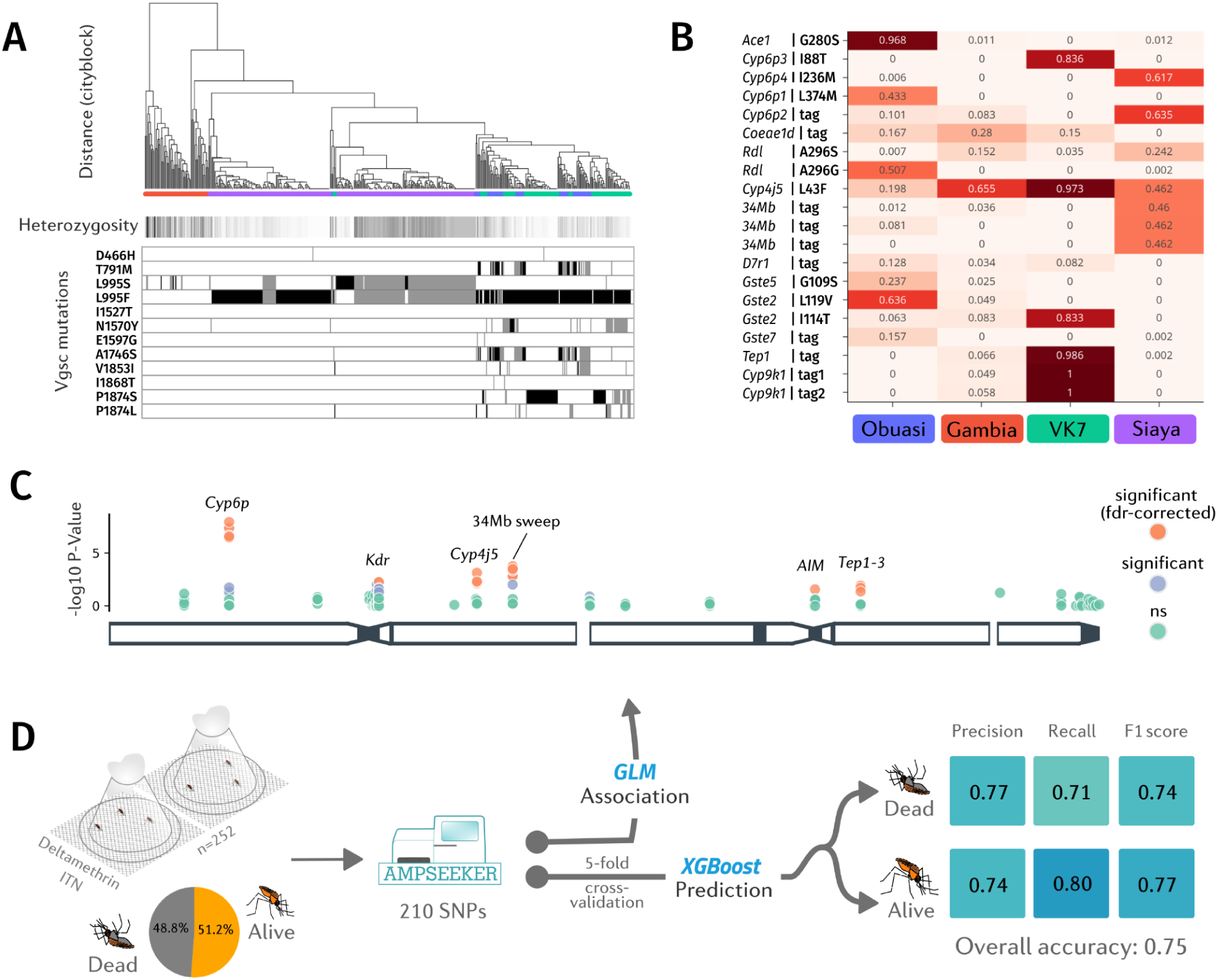
Genomic surveillance of insecticide resistance with Ag-vampIR and AmpSeeker. A) Diplotype clustering at the Voltage-gated sodium channel. We calculate pairwise distance between diplotypes at all *Vgsc* sites. Each column in the figure is a diplotype ordered by the dendrogram by hierarchical clustering, using genetic distance based on city-block (Manhattan) distance and complete linkage. The leaves of the dendrogram are colored by the species of the individual to which they belong. Underneath the dendrogram, the heterozygosity (calculated as an average over all SNPs in the *Vgsc* locus) is displayed as horizontal bars. Clusters with low sample heterozygosity or inter-sample genetic distances of 0 are indicative of a selective sweep. Amino acid variation is displayed below the dendrogram - black indicates homozygosity and grey heterozygosity. B) Allele frequencies at other known insecticide-resistance SNPs targeted by Ag-vampIR. C) A Manhattan plot of the amplicon-wide association study using a generalised linear model. Siaya mosquitoes were exposed to deltamethrin-treated nets (PermaNet 2.0) in extended 40 minute WHO cone assays. D) A schematic of the approach to predicting the insecticide resistance phenotype from WHO cone assays, highlighting the Precision, Recall, F1-score and overall accuracy of the XGBoost algorithm in both prediction classes (Dead & Alive).

### Allele frequencies at resistance-associated SNPs

We next surveyed other resistance-associated mutations targeted by Ag-vampIR (Figure 3B). The *Ace1-*G280S mutation, known to confer resistance to organophosphates and carbamates, was found close to fixation in *An. gambiae* from Obuasi (97% frequency), but was completely absent in VK7, with very few resistance alleles found in the Gambia (1) or Siaya (6). The two *Rdl* mutations A296G and A296S, which confer resistance to dieldrin, remain at moderate frequencies in sub-Saharan Africa, despite a ban on dieldrin since 1974. We found A296S at 24.2% in Siaya, but rare or absent in other cohorts, whereas A296G was at 50.7% frequency in the Obuasi cohort, but absent in other cohorts.

We also monitor metabolic resistance variants, many of which were selected to track selective sweeps revealed by phase 1 of the Ag1000g (Supplementary Figure 9). Eight amplicons target the Cyp6p locus, and we find the *Cyp6p3-*I88T mutation at high (83.6%) frequency in VK7. The *Cyp6p4-*I236M mutation, which is associated with a gene duplication in East Africa ^37^, is at 61.7% frequency in Siaya. The *Cyp6p1-*L374M mutation at high frequency in Obuasi, tags an ongoing selective sweep found in *An. gambiae* from West Africa, originally identified in samples from Cameroon and Burkina Faso. Two mutations at *Cyp9k1* that tag a selective sweep are fixed in VK7 (100%), whilst again present at low frequencies in The Gambia (5.2%, 7.7%). Consistent with this, these SNPs tag a selective sweep at *Cyp9k1* originally found in *An. coluzzii* from Burkina Faso ^13^. At the *Gste* gene cluster, members of which have been associated with resistance to DDT, organophosphates, and pyrethroids, the *Gste2-*I114T mutation is at 83.1% frequency in VK7, but at low frequencies in the Obuasi (6.7%) and Gambia (6.8%) cohorts and absent in Siaya. The *Gste2-*L119V mutation is at 61.9% frequency in Obuasi, and 5.4% in The Gambia. The low frequency of resistance mutations in The Gambia is likely due to the dominance of *An. arabiensis* in the cohort.

The *Cyp4j5-*L43F mutation, which has repeatedly been associated with resistance to pyrethroids in East Africa ^38,39^, is found in all cohorts at moderate frequencies, 19.8% in Obuasi, 65.5% in the Gambia, 97.3% in VK7, and 46.2% in Siaya. In East Africa (and therefore Siaya), this mutation is at very high frequency on the 2La inversion karyotype, and is therefore acting as a tagging SNP for the inversion. The same may not be true in West Africa, where it can reach higher frequencies and has not been associated with pyrethroid resistance. Three tagging SNPs tag a selective sweep at approximately 34Mb on the 2L chromosomal arm, again found on the 2La inversion karyotype. This sweep is ongoing in *An. gambiae* from East Africa, and appears to be at 46% frequency in Siaya, whilst absent from other cohorts.

We also find the TEP1 *Plasmodium*-refractoriness allele fixed in VK7 (100%), and at low frequencies in The Gambia (8.3%). AmpSeeker also reports allele frequencies of mutations not specifically targeted by Ag-vampIR, which we provide as an exercise for the reader (https://sanjaynagi.github.io/agvampir002-results/notebooks/allele-frequencies.html).

### Genotype:phenotype association study

The much lower cost of amplicon sequencing compared to whole-genome sequencing enables association studies with larger sample sizes. The samples from the Siaya cohort were phenotyped against new PermaNet 2.0 deltamethrin-impregnated nets and we tested whether any variants genotyped by Ag-vampIR were associated with mosquito survival. For each SNP, we built a binomial GLM with the SNP as the predictor and phenotype as the response variable, followed by false discovery rate control to correct for multiple testing. For this analysis, we utilised any SNPs present on Ag-vampIR amplicons, after filtering, this resulted in 210 SNPs. The full results of the association study are in Supplementary Data 2 and are visualised as a manhattan plot in Figure 3C.

After false-discovery rate control, thirty-seven SNPs were significantly associated with survival following exposure to PermaNet 2.0 nets, corresponding to seven independent mutations. Thirteen SNPs in the Cyp6p gene cluster, including *Cyp6p4-*I236M, were positively and strongly associated with mosquito survival. All thirteen showed highly similar odds ratios ranging between 3.16 and 3.65. The presence of *Cyp6p4-*I236M suggests that Siaya harbors the triple-mutant haplotype described in Njoroge *et al*., ^37^, and the twelve remaining SNPs are therefore all SNPs found on this mutant haplotype and are tagging *Cyp6aa1-*Dup1. At the *Vgsc*, the 995F mutation was significantly positively associated with survival (OR=1.78, 95% CIs=1.18-2.68), whereas 995S was negatively associated (OR=0.56, 95% CIs=0.37-0.84). These data are consistent with the literature, suggesting that 995F confers greater resistance to deltamethrin, and that 995F is currently replacing 995S in East Africa ^17,35^.

Four SNPs at *Cyp4j5* were associated with resistance, including the non-synonymous mutations *Cyp4j5-*A47P (OR=1.78, 1.16-2.72), *Cyp4j5-*L43F (OR=2.25, 1.40-3.60), and *Cyp4j5*-T30W (OR=1.94, 1.22-3.07). Fourteen SNPs at the 34Mb sweep locus are associated with survival; all with similar odds ratios ranging from 2.10 to 2.48. In Ag1000g data, there is only a single selective sweep at this locus, and so these fourteen SNPs are likely to be tagging this selective sweep. This is the first evidence that this haplotype confers resistance to insecticides used in vector control. A SNP (3L:380924, C), which is found on the same amplicon of an ancestry informative marker (3L:380974, C), is negatively associated with survival (OR=0.56, 0.37-0.83). The SNP is located in an intron of AGAP029563, a gene without a description or any functional annotation. Three intergenic SNPs at the TEP locus, previously associated with *Plasmodium* refractoriness, were also associated with mosquito survival, in agreement with recent GWAS results suggesting that this locus may also be important for insecticide resistance^14^. One SNP (3L:11214067, G) was positively associated (OR=1.96, 1.20-3.21), whilst two others, likely to be found on a competing haplotype, were strongly negatively associated with survival with odds ratios of 0.29 (0.14-0.63) and 0.33 (0.16-0.68).

### Predicting bioassay survival with machine learning

Given the number of loci associated with bioassay survival to insecticide-treated nets, we hypothesised that it may be possible to build models using machine learning that are predictive of bioassay mortality. We trained an XGBoost model on the 252 phenotyped Siaya samples at the 210 genotyped SNPs using five-fold cross-validation, to predict the mortality of individual mosquitoes. Figure 3D shows a schematic of the phenotypic prediction using XGBoost.

The model was able to predict with 75% accuracy (weighted F1-score of 0.75) the classification of individual mosquitoes as either dead or alive in the bioassay. Cross-validation demonstrated consistent performance across different data splits (CV score: 0.754 ± 0.116). Analysis of feature importance revealed several key genetic variants driving the predictions, with the most influential SNPs located at 2L:28502739 (*Cyp6p* cluster), 3L:11208428 (*Tep* cluster), and 2L:34100775 (34Mb sweep), in agreement with the earlier association study. The model showed balanced performance between predicting both dead (precision: 0.77, recall: 0.71) and alive phenotypes (precision: 0.74, recall: 0.80), suggesting robust predictive capability across both outcomes. This substantially outperformed random baseline models (0.37-0.49), demonstrating that the genetic variants captured meaningful signals predictive of insecticide resistance.

### Cost-effectiveness of Ag-vampIR

In Supplementary Table 6, we estimate costs of reagents and kits used in the Ag-vampIR protocol. The estimated per sample cost, based on 500 samples run on a MiSeq reagent kit V2, is $9.23, which equates to ∼$0.10 for each of the ninety loci targeted. For comparison, whole-genome sequencing is estimated to cost over $100 dollars per sample, whilst an LNA probe-based qPCR assay would cost an estimated ∼$2.475 per locus genotyped. These costs include performing DNA extractions with a column-based proprietary kit (Nexttec), the most expensive part of the protocol, which could feasibly be removed for a simpler, cheaper extraction ^40^.

## Discussion

We have developed an integrated lab protocol and computational workflow to perform effective, targeted, end to end genomic surveillance of *An. gambiae s.l*. We show that Ag-vampIR and AmpSeeker can rapidly generate accurate and readily-interpretable data at an affordable cost, opening up novel areas of research and enabling widespread genomic surveillance.

Overall, the cost-effectiveness of the platform allows genomic surveillance on a much larger scale, in comparison to whole-genome sequencing ^14,41^. We demonstrate how this can be particularly useful in performing genotype:phenotype association studies, boosting power to detect variants involved in the insecticide-resistant (or any other) phenotype. The association study with the Siaya colony yielded important insights into pyrethroid resistance mechanisms, identifying seven independent variants significantly associated with survival against PermaNet 2.0 nets. The study’s findings validated several known resistance mechanisms while uncovering new associations, for example, at the previously uncharacterized 34Mb sweep locus.

The demonstration that Ag-vampIR genotypes alone could predict WHO cone bioassay mortality with good accuracy is particularly promising. The XGBoost model’s performance (weighted F1-score of 0.75) suggests that genetic data could potentially serve as a proxy for phenotypic resistance testing, though further validation across different populations and bioassay protocols is necessary. Whilst strong predictive power for organophosphate and carbamate resistance has been demonstrated for the Ace-1 G280S mutation ^16,42^, high prediction capacity for pyrethroid resistance is impressive given the multivariate nature of resistance mechanisms in contemporary African populations of *An. gambiae* ^14,43^. This capability could transform resistance monitoring by enabling rapid, cost-effective screening of vector populations, allowing control programmes to make data-driven decisions about insecticide deployment. Whilst phenotypic testing will still play a role, this predictive approach could enable more frequent and widespread resistance surveillance than is currently feasible with bioassays.

Our approach also proved highly effective for exploring population structure and ancestry in our samples. By targeting ancestry informative markers (AIMs), we were able to both distinguish between *An. gambiae* and *An. coluzzii* and accurately identify other sibling species including *An. arabiensis* and *An. melas*. The high accuracy of our machine learning approaches in species classification (F1 scores >0.99) demonstrates the discriminatory power of these markers, a potentially surprising finding, given that they were intended to differentiate only *An. gambiae* and *An. coluzzii*.

Robust and standardised tools that facilitate the rapid and reproducible analysis of genomic data are required for effective genomic surveillance. Whilst bioinformatic capacity in sub-Saharan Africa is increasing, there remains a paucity of standardised analysis methods for genomic data, outside of the Vector Observatory^44^. In developing AmpSeeker, we provide an all-purpose tool for scalable and reproducible analysis of Illumina amplicon sequencing data. AmpSeeker’s novelty is in its generation of a local webpage of results, allowing the user to explore the data interactively, and enabling the rapid, interpretable dissemination of results to partners and stakeholders alike. The interactive nature of the results, which includes descriptive, explanatory text, we hope facilitates an understanding of complex genomic data, potentially bridging the gap between bioinformaticians and field researchers or policymakers. AmpSeeker is open-source and written in Python, a mature and popular programming language for bioinformatics, increasing the likelihood of long-term sustainability and ease of maintenance. Its open-source nature aligns with the principles of open science, promoting transparency and collaboration. We are currently delivering workshops and collaborating with research groups across sub-Saharan Africa to facilitate its implementation. We are also expanding the utility of AmpSeeker to amplicon panels from other organisms. Given its applicability to any amplicon sequencing data, and the ease to which analysis of other panels can be incorporated, we expect AmpSeeker to prove useful to the research community beyond vector amplicon panels. We strongly encourage the vector and genomics communities to get involved in AmpSeeker’s development, which we hope will foster community ownership and potentially drive wider adoption.

Centralised genome sequencing efforts such as the Ag1000g and Pf7, have advanced the field, generating large amounts of high-quality genomic data and revealing major evolutionary and public health insights ^45,46^. Routine genomic surveillance, however, would be more effective if decentralised and conducted in-country, to enable the rapid turnaround from sample collection to results, to cultivate research ownership, and to facilitate effective communication with NMCPs and malaria stakeholders. The development of complete workflows for genomic surveillance, alongside ongoing efforts in building genomics capacity, will be pivotal to this decentralisation process. However, major challenges remain - for example, the cost of obtaining reagents for sequencing (and indeed, other molecular biology applications), in sub-Saharan Africa (SSA) remains unacceptably high; Although we show that our protocol is a cost-effective route to high-throughput genotyping, the cost in SSA is likely to be two or three times higher. Existing genomic infrastructure must be supported and expanded further with sustained investment in both equipment and expertise ^47^. Critical attention must be paid to establishing reliable supply chains for reagents and consumables, while developing regional centers of excellence that can provide training and technical support. Importantly, in the case of amplicon sequencing, analysis can be conducted on any standard bioinformatic laptop running Linux or Mac, eliminating the need for specialized computing infrastructure. Additionally, mechanisms for data sharing and standardization across countries need to be established to ensure that decentralized efforts contribute to a coherent understanding of vector populations across the continent.

A fundamental limitation of any targeted sequencing approach is the inherent bias towards known variants and genomic regions of interest. While our panel captures many known resistance mechanisms in *An. gambiae s.l*, novel mutations arising outside our targeted amplicon regions will inevitably be missed. This is particularly relevant given the remarkable adaptability of *Anopheles* populations and their demonstrated capacity to evolve new resistance mechanisms under insecticide pressure ^13^. Efforts are underway to generate a web resource of selection signals in the Vector Observatory (https://anopheles-genomic-surveillance.github.io/selection-atlas/home-page.html), with the aim of building a database of haplotypes under putative selection, which we will use to update the Ag-vampIR panel. A second limitation of the approach is its limited ability to detect large structural variants which could mean that CNVs that confer metabolic resistance are missed, although they may be tagged by tagging SNPs. The modular nature of AmpSeeker and our panel design allows for updating the panel. This will be vital for ensuring long-term success of the platform. Using AnoPrimer, we can design primers for additional loci with ease whilst avoiding regions where genetic variants are at appreciable frequency in wild populations *An. gambiae s.l*^48^. Future versions of the panel could incorporate recently discovered markers of resistance ^19,49^ and tagging SNPs for major chromosomal inversions.

## Conclusion

The development of complete workflows for genomic surveillance, alongside ongoing efforts in building genomics capacity, will be central to effective malaria vector control. Through our approach, we provide an integrated solution for targeted genomic surveillance of insecticide resistance and species composition in the *An. gambiae* species complex. The platform’s ability to track multiple resistance mechanisms and generate reproducible and interpretable results makes it a valuable tool for evidence-based decision making. As insecticide resistance continues to threaten malaria control efforts, accessible and standardized approaches to surveillance will become increasingly critical. By providing open-source tools and protocols, we hope to contribute to the development of sustainable, decentralized genomic surveillance capabilities across disease-endemic regions.

## Materials and Methods

### Sample collections and bioassay testing

Samples from four cohorts were collected and sequenced with the Ag-vampIR protocol (Supplementary Table 1), that spanned different species and geographies across sub-Saharan Africa. This included two laboratory colonies, Siaya and VK7, and two field cohorts from Obuasi, Ghana, and the eastern Upper River Region, The Gambia. Siaya (*An. gambiae)* were established in 2023 from Siaya county, Western Kenya, and VK7 (*An. coluzzii)* in 2014 from Valle de Kou, Burkina Faso. Obuasi samples were collected in 2017 with indoor resting aspiration by Prokopak aspirators, whilst the Gambia samples were sampled as part of the SANTE study with CDC miniature light traps ^50^.

For the association study, we exposed 3-5 day old female mosquitoes from the Siaya colony to new, unopened PermaNet 2.0 (deltamethrin) LLINs in extended WHO cone test bioassays^51^. The WHO cone test exposure time was extended from 3 to 40 minutes to induce close to 50% mortality. The details of each specimen involved in the study are specified in Supplementary Data 3.

### Ag-vampIR primer and panel design

Details of the design of Ag-vampIR are found in Supplementary Text 2. In brief, primer design for all amplicons except Agam-79 (*Vgsc-*V402L), Agam-80 (I1527T) and *Dsx* was performed using an in-house modified version of mprimer^52^. The program modifications speed up the process of multiplex PCR design without any effect on the resultant primer design output itself. The V402L and I1527T amplicons were added later, designed with AnoPrimer, a primer design tool that considers the presence of genetic variation from the Ag1000g in primer binding sites^48^. For each input file, mprimer was run using the following parameters: primer min_gc20, primer opt_gc 50, primer max_gc 80, primer min_tm 50, primeropt_tm 58, primer max_tm 68, product_min_size 190 bp, product_max_size 250 bp, penalty allowance 10.

To prepare samples for Illumina sequencing with sample barcoding (detailed below), we modified each primer by adding standard Illumina adapter sequences. The forward adapter sequence 5′-ACACTCTTTCCCTACACGACGCTCTTCCGATCT- was added to the locus-specific forward primer, while the reverse adapter sequence 5′-TCGGCATTCCTGCTGAACCGCTCTTCCGATCT- was added to the locus-specific reverse primer. Additionally, to reduce primer dimer formation, we replaced the second-to-last base with a 2’-O-Methyl base, following the approach described by McKerrell et al. ^53^. Primer sequences are detailed in Supplementary Data 1.

### Laboratory protocol and sequencing

DNA was extracted from individual mosquitoes using Qiagen DNeasy extraction kits. For convenience, we did not quantify or normalise DNA concentrations prior to the Ag-vampIR protocol, however, this would be expected to result in more even coverage between sequencing plates. Full protocols are published in Jacob et al.,^22^, with modifications below. In brief, an initial locus-specific PCR (PCR1) is run on each individual DNA sample, using a primer pool of all primers at 10μm. PCR1 is immediately followed by PCR2, in which PCR1 products are added to plates containing lyophilised i5 and i7 sample-specific barcodes and Illumina adaptors. In PCR2, these barcodes are added to the ends of the locus-specific amplicons, and permit hybridisation to the Illumina flow cell. After PCR2, each plate is pooled together, followed by an AMPure XP bead clean up and fragment analysis with an Agilent Tapestation or BioAnalyzer. qPCR is then used to determine the concentration of the final libraries, which are then diluted and pooled and prepared for sequencing according to Illumina protocols. We sequenced the libraries on an Illumina MiSeq with a MiSeq reagent kit V2, which provides 150 Bp reads and a total output yield of ∼5 Gb.

### Bioinformatic analysis

Full details of the AmpSeeker pipeline are designated in Supplementary Text 3. AmpSeeker is written in snakemake^20^, according to the best practices. In brief, AmpSeeker converts Illumina BCL files to fastq with BCL-Convert (v4.3.6). Reads are then trimmed and assessed for quality with fastp ^54^, and the quality of index reads is assessed separately with fastqc^55^. Reads are then aligned to the reference genome with bwa mem^23^, with indexing performed by samtools^56^. Coverage is assessed with mosdepth^57^, and alignments visualised with an IGV notebook^58^. Variants are then called with bcftools mpileup^24^ and annotated with snpEff ^59^.

For analysis of the resulting variant calls, AmpSeeker utilises a series of custom-built Jupyter notebooks written in Python. Notebooks are executed by snakemake using the python tool papermill, which allows parameters to be passed to the notebook prior to its execution. For genomic analysis, we use scikit-allel v1.3.7^60^, whilst all visualisation is performed with Plotly ^61^ for its interactive plotting capabilities. Executed jupyter notebooks, containing data and interactive plots, are then combined with Jupyter Book^62^ to generate a HTML web page containing all data and results.

### Ancestry analysis and taxon assignment

We employed three complementary approaches to assign taxonomic status to our samples: Ancestry Informative Markers (AIMs), and two machine learning methods applied to principal components (PCA), and genotypes directly (XGBoost). Ancestry informative markers to distinguish *An. gambiae* from *An. coluzzii* were ascertained from Miles *et al.,*^13^ which used a subset of SNPs from Neafsey *et al.,*^63^, with samples from Mali. We used the AIMs to paint genotypes in our samples and to assign a putative taxon to samples. To check the reliability of our AIMs against *An. gambiae* from East Africa, we checked concordance of each AIM with any public samples from Uganda, Kenya, or Tanzania which were assigned as *An. gambiae* by the vector observatory pipelines^44,46^. For taxon assignment, we only use AIMs found on chromosome 3 and the X chromosome, to avoid known regions of adaptive introgression on chromosome 2 ^30^. We called samples with an average AIM genotype of less than 0.5 as *An. gambiae,* between 0.5 and 1.5 uncertain, and above 1.5 called as *An. coluzzii*.

We built an extreme gradient boosting (XGBoost) model, using all Ag-vampIR SNPs as input, by training a model on 7363 curated, QC-controlled, whole-genome sequenced individuals from the Ag1000g, and used SNP data from five taxa in the complex: *An. gambiae, An. coluzzii, An. arabiensis, An. melas,* and the Pwani and Bissau molecular forms (Supplementary Table 2). We removed species with less than 10 samples (*An. quadrianulatus*, *An. merus* and *An. fontenillei*). The model was implemented using the XGBoost framework in Python, using 5-fold cross-validation. Feature selection was performed using a mean threshold criterion to identify informative AIMs, and hyperparameter optimization was conducted through a randomised cross-validation search. For the XGBoost model, we optimized parameters including learning rate (0.01-0.3), maximum tree depth (3-9), number of estimators (100-300), minimum child weight (1-5), and subsample ratio (0.6-1.0). The XGBoost model was highly accurate on Ag1000g data (Supplementary Table 3). Model performance was evaluated using precision, recall, and F1-scores, with confusion matrices generated to assess classification accuracy across all species.

We also performed Principal Component Analysis (PCA) on combined data from our amplicon samples and 7,363 whole-genome sequenced reference samples from the Vector Observatory. After removing invariant sites, we conducted PCA on the remaining variable sites and used the first four principal components for classification. To automatically assign taxon status, we implemented a Support Vector Machine (SVM) classifier with a probability threshold of 0.8 to assign taxonomic status. Samples with classification probability below this threshold were marked as “unassigned.” This approach allowed us to visualize the genetic relationships between our samples and reference populations in reduced dimensional space while providing statistical confidence for taxonomic assignments. For each sample, we assigned the taxonomic status that was predicted by at least two of the three methods. In cases where no consensus was reached (i.e., all three methods predicted different taxa), samples were labeled as “unassigned.“

### Population structure

For population structure analyses, we utilised variants present on any amplicon, not just those SNPs specifically targeted by the Ag-vampIR panel. Performing population structure analyses with resistance loci comes with the caveat that in many cases these loci will be non-neutral, given that they may be under selection, whilst ancestry informative markers are expected to segregate by species. After filtering out any SNPs that were invariant or missing in more than 5% of samples, 2389 SNPs remained. AmpSeeker both performs principal components analysis (PCA) and builds Neighbour-joining trees (NJT) generating interactive plots, coloured by user-selected metadata.

### Kdr analysis and Diplotype clustering

Diplotype clustering of the Voltage-gated sodium channel was performed as in Nagi et al.,^19^, and is as follows: We extracted diplotypes from the Vgsc gene region and performed complete-linkage hierarchical clustering analysis. Diplotypes, which can also be called multi-locus genotypes, represent sequences of diploid genotypes. We first converted diplotypes into allele count arrays, where each site is represented by an array of four elements corresponding to the possible alleles. For each site, we calculated the pairwise differences between these allele count arrays for all pairs of individuals. The clustering was performed using city-block (Manhattan) distance as the distance metric with complete linkage. We also identified the non-synonymous variants present on each diplotype and plotted this information alongside sample heterozygosity across the region.

We included heterozygosity in the diplotype clustering plot, to permit easy identification of diplotype clusters homozygous for a selective sweep, as opposed to diplotype clusters which contain two separate haplotypes under selection. To calculate individual-level heterozygosity, we used the scikit-allel v1.3.7 implementation (allel.GenotypeArray.is_het). For a given diplotype region, we calculated the number of heterozygote calls and then divided by the number of called alleles to get a single value of heterozygosity for the region for each individual sample.

### Genotype:phenotype associations

For the Siaya colony genotype:phenotype association tests, we loaded SNPs from all amplicons, removing invariant, highly missing (>20% of calls), or low minor allele frequency SNPs (<2%). We then split multi-allelic calls, and converted genotypes to the number of alternate calls. For each SNP, we ran a binomial generalised linear model (GLM) with logit link function using the number of alternate calls at the SNP of interest as a predictor variable and the phenotype (dead or alive in WHO cone test) the response variable. We then applied FDR correction to the resulting p-values with statsmodels^64^.

### Predicting bioassay mortality with an XGBoost machine learning model

We developed a machine learning approach using XGBoost (eXtreme Gradient Boosting). We used data from the Siaya colony mosquitoes that had been exposed to PermaNet 2.0 nets in extended WHO cone bioassays. For features, we utilized 210 quality-controlled SNPs that passed filtering for variance and missingness from the Ag-vampIR amplicons. We used 5-fold cross-validation to assess model performance, ensuring consistent evaluation across different data partitions.

We optimized hyperparameters through a cross-validation grid search (scikit-learn’s GridSearchCV), exploring ranges for key parameters including learning rate (<u>0.01</u>, 0.1, 0.3), maximum tree depth (3, 7, <u>12</u>), number of estimators (<u>100</u>, 500), minimum child weight (<u>1</u>, 5), gamma (0, <u>0.3</u>), subsample ratio (0.7, <u>1</u>), reg_alpha (<u>0</u>, 1) and reg_lambda (<u>1</u>,4). The final best model parameters are underlined.

## Data availability

The results book for this analysis is located at sanjaynagi.github.io/agvampir002-results/, which we encourage the reader to explore. Read data is deposited in the SRA (BioProject accession PRJNA1207724).

## Author contributions

MJD, SCN, TN, JK, DW and ERL conceptualised and designed the study. SCN, ERL, CGJ, TP, AVH, CH, KR, AJ, ScG, NP, CA, SR and KR contributed to the development of the Ag-vampIR panel and laboratory protocol. SCN, TM, EL, ERL, AHK and AM contributed to the development of the AmpSeeker pipeline. FA, CY, ML and SS carried out DNA extractions and performed sequencing. MJD, VJS, SoG, TN, JK secured the funding to support the sequencing and analysis. HN, JE, AEY and JW provided samples. SCN led the data analysis with contributions from ERL, FA, SS and AM. SCN wrote the manuscript with contributions from all co-authors.

## Acknowledgements

The authors would like to extend our deepest gratitude to the late Professor Dominic Kwiatkowski, whose pioneering contributions to the field of malaria genomic epidemiology paved the way for this research. We would also like to acknowledge the MalariaGEN laboratory teams at the University of Oxford and the Wellcome Sanger Institute for their contributions to the laboratory protocol, and to the SANTE trial team for mosquito collections. We thank Petra Korlevic for her illustrative logo for the Ag-vampIR panel.

This work was supported by the National Institute of Allergy and Infectious Diseases (NIAID R01-AI116811 to M.J.D. and D.W.) and the Medical Research Council (MR/T001070/1 to M.J.D., D.W., and E.R.L., MR/P02520X/1 to M.J.D. and D.W., MR/W002159/1 to T.N.). The latter grant is a UK-funded award and is part of the EDCTP2 program supported by the European Union. M.J.D. was supported by a Royal Society Wolfson Fellowship (RSWF\FT \180003). SCN was funded by an MRC CASE studentship (MR/R015678/1). VJS is supported by the Bill and Melinda Gates Foundation (INV-068808). This study/project is funded by the NIHR [NIHR Global Health Research Group on Establishing Regional Hubs for Genomic Surveillance in West Africa (NIHR 134717)]. The views expressed are those of the authors and not necessarily those of the NIHR or the Department of Health and Social Care.^14^

