## Supplementary Figures for "Targeted genomic surveillance of insecticide resistance in African malaria vectors"

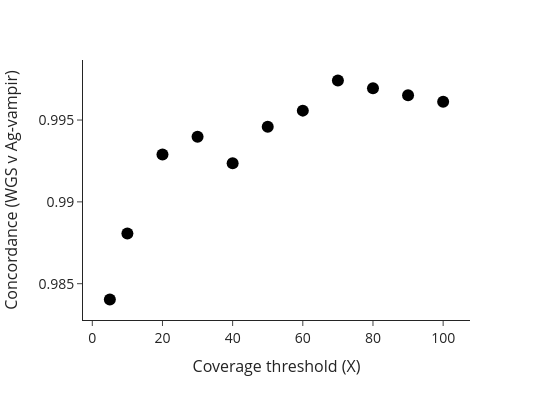

**Supplementary Figure 1.** Concordance of Ag-vampIR variant calls with whole-genome sequencing (WGS) data. Concordance was evaluated across 11 variant read depth thresholds using 40 individuals from Obuasi, Ghana, previously sequenced to 30x coverage as part of the Ag1000g project. High concordance (>99.3%) was observed at read depths of 20x and above, demonstrating the reliability of the Ag-vampIR-AmpSeeker platform.

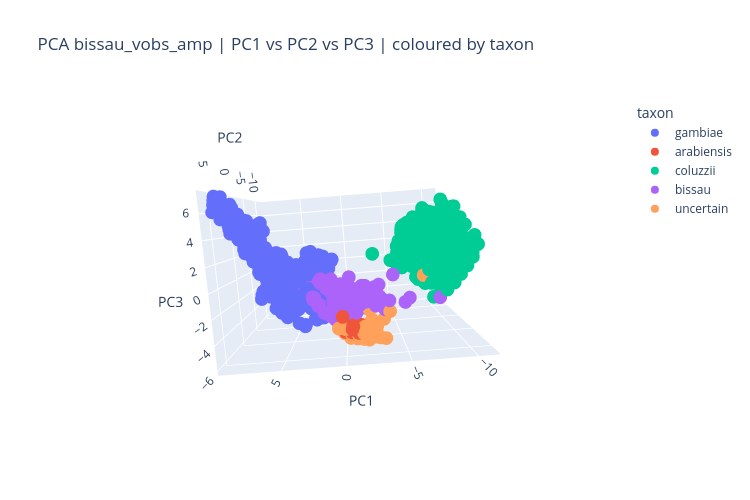

**Supplementary Figure 2**. Principal components analysis (PCA) of AmpSeq samples from The Gambia and Ag1000g samples from across sub-Saharan Africa. PCA was performed using SNPs from the Ag-vampIR panel.

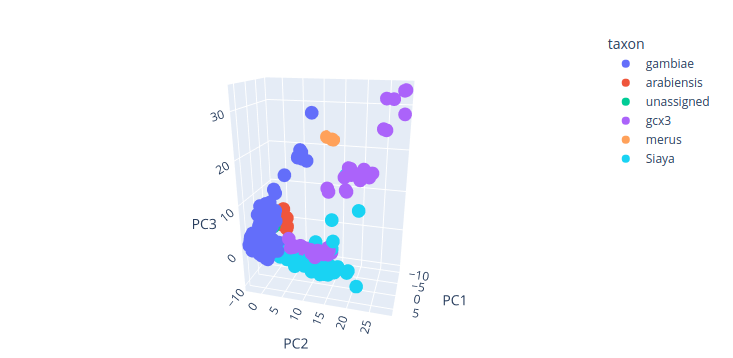

**Supplementary Figure 3**. 3D Principal components analysis (PCA) of whole-genome sequenced mosquitoes from the Ag1000g dataset and Siaya colony samples. PCA was performed using SNPs targeted by the Ag-vampIR panel. Siaya samples cluster near the An. gambiae cluster and adjacent to the Pwani molecular form (Gcx3), suggesting potential shared ancestry.

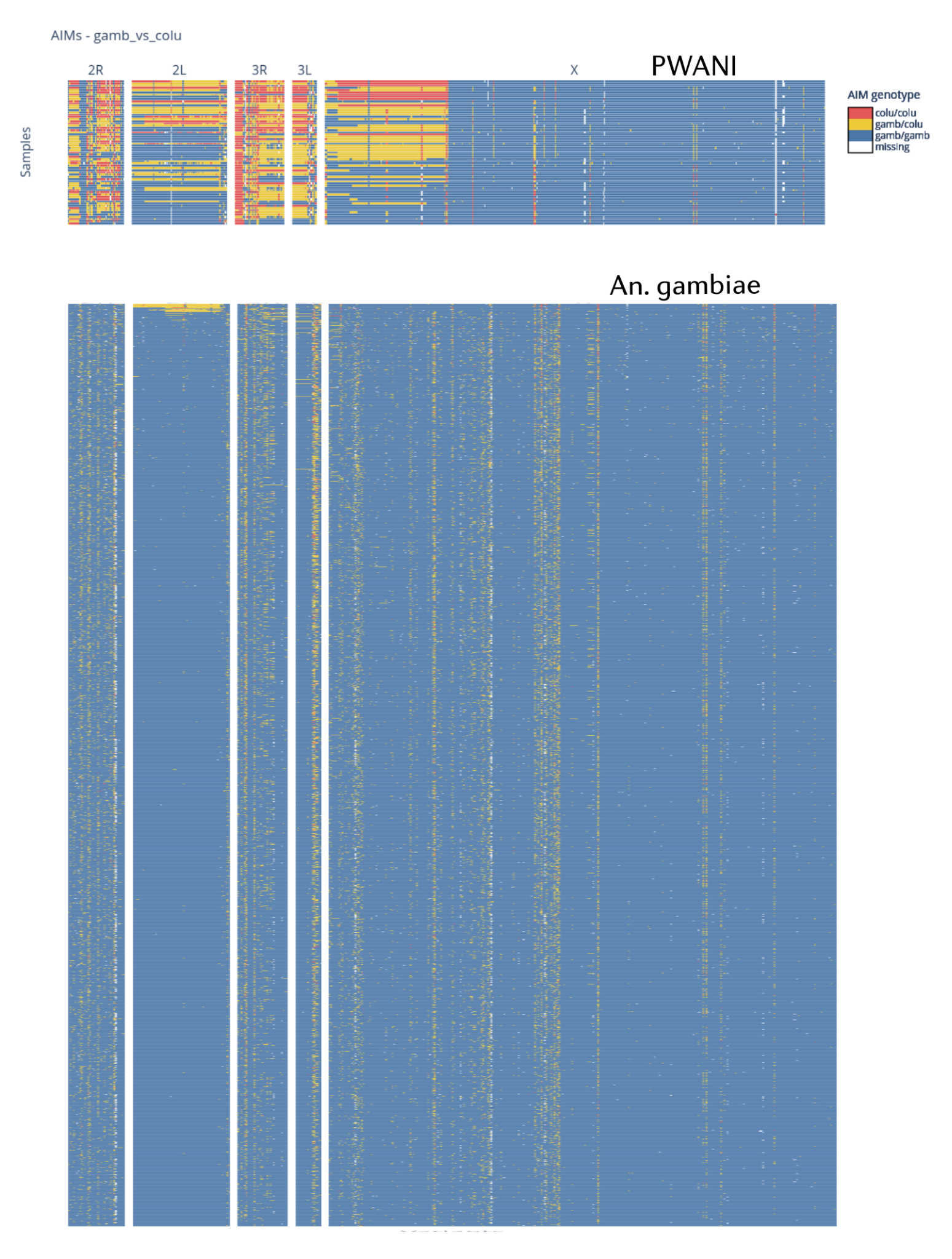

**Supplementary Figure 4.** Ancestry informative marker (AIM) genotypes for Pwani molecular form (Gcx3) and An. gambiae samples. AIM patterns on autosomes for Pwani samples appear mixed, while An. gambiae samples consistently show An. gambiae-like AIMs, highlighting the distinct ancestry profiles of these taxa.

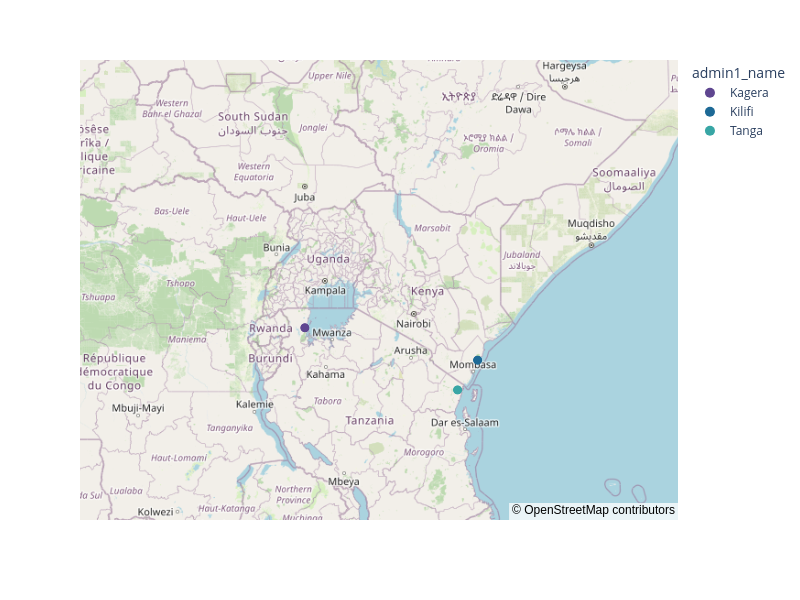

**Supplementary Figure 5:** Geographic distribution of the Pwani molecular form (Gcx3) in East Africa. A single Pwani individual from the Ag1000g dataset was collected in Muleba, Tanzania, on the western coast of Lake Victoria in 2015, suggesting that this taxon may also exist in inland areas.

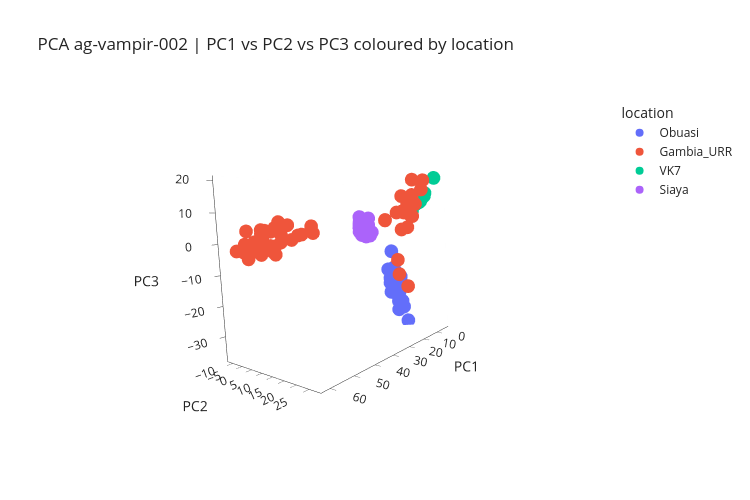

**Supplementary Figure 6:** 3D Principal components analysis (PCA) of Ag-vampIR samples, colored by cohort. PCA reveals five distinct clusters corresponding to different cohorts, including the Bissau molecular form, Siaya colony, Obuasi An. gambiae, and two An. coluzzii clusters (VK7 and a subset of Gambian samples).

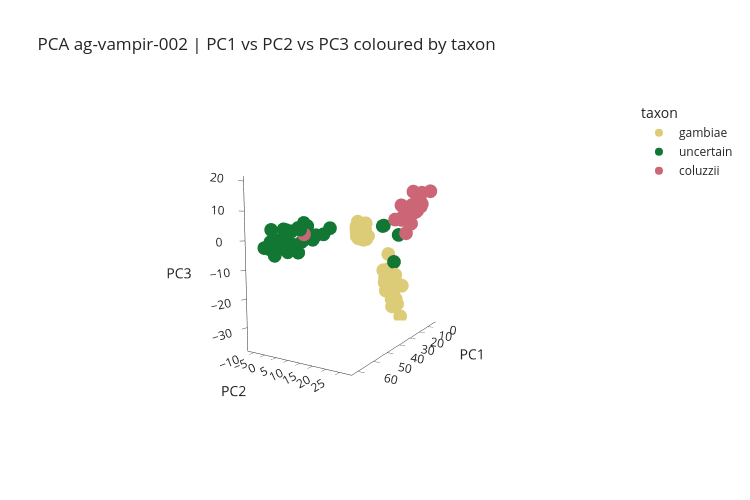

**Supplementary Figure 7:** 3D Principal components analysis (PCA) of samples, colored by taxon assignment. Taxon assignments were based on ancestry informative markers (AIMs) targeted by the Ag-vampIR panel.

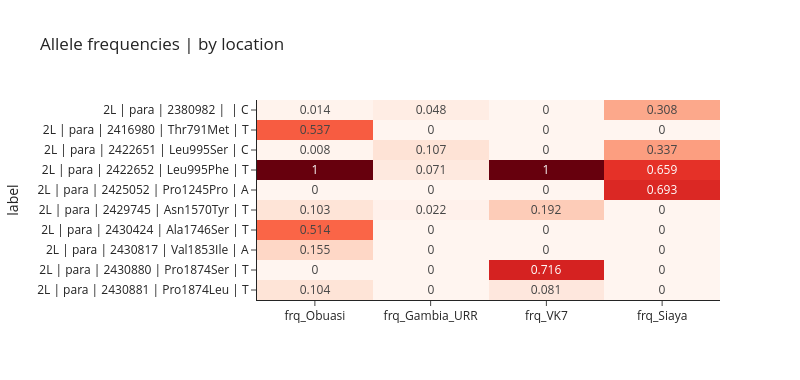

**Supplementary Figure 8:** Allele frequencies of known resistance-associated SNPs in the Voltage-gated sodium channel (Vgsc). Frequencies were calculated using SNPs targeted by the Ag-vampIR panel, with tagging SNPs used to infer Kdr haplotypes.

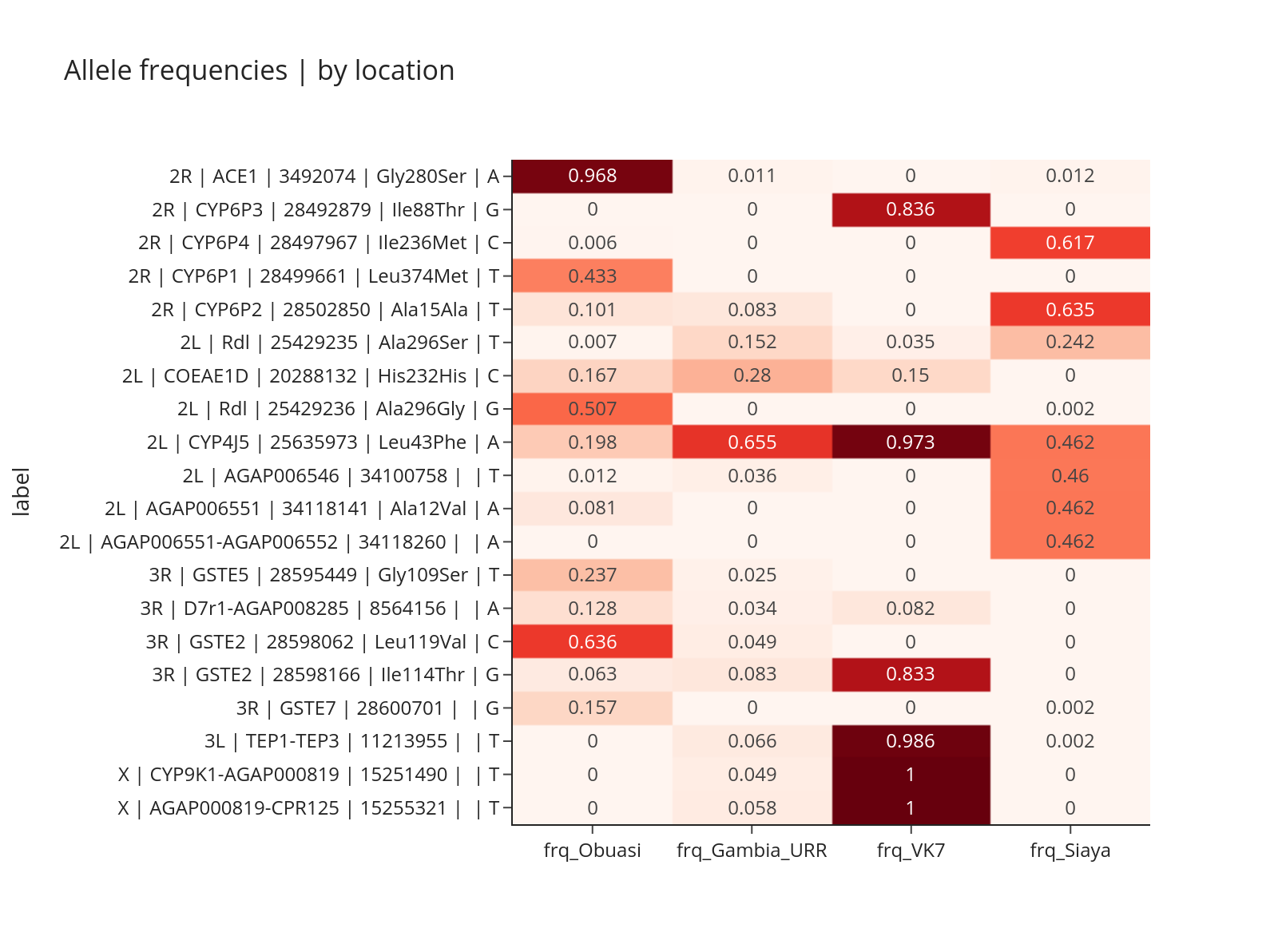

**Supplementary Figure 9:** Allele frequencies of known resistance-associated SNPs excluding the Voltage-gated sodium channel (Vgsc). Frequencies were calculated using SNPs targeted by the Ag-vampIR panel, with tagging SNPs used to infer Kdr haplotypes.
