## Supplementary Text 1 for "Targeted genomic surveillance of insecticide resistance in African malaria vectors"

### **Supplementary Text 1 - Lab protocols**

The lab protocols are published in:

Christopher G Jacob, Nguyen Thuy-Nhien, …, Dominic P Kwiatkowski, Olivo Miotto (2021) **Genetic surveillance in the Greater Mekong subregion and South Asia to support malaria control and elimination** eLife 10:e62997

With the following modifications for application to An. gambiae:

| **Comparison of Pf and Ag-vampIR protocol** | | | |
| --- | --- | --- | --- |
| **Protocol** | **Feature** | **Pf** | **Ag-vampIR** |
| Overall protocol | Number of panels | 3 | 1 |
| PEP/sWGA | Whole genome amplification | sWGA | PEP |
|  | Conc for T&B samples | 1.33ng/ul | 1ng/ul |
| Library prep  (T&B / Operational) | PCR_1 thermocycler parameters | 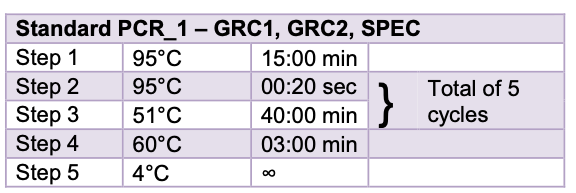  Annealing temp = 51C | 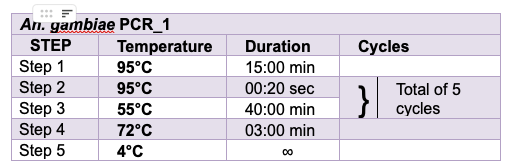  Annealing temp = 55C |
| Library prep  (T&B / Operational) | PCR_2 thermocycler parameters | 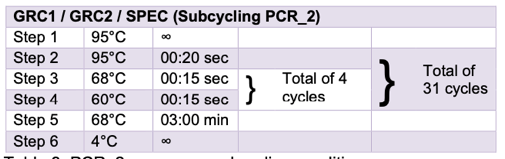  Temp cycles between 68C and 60C for 15s each | 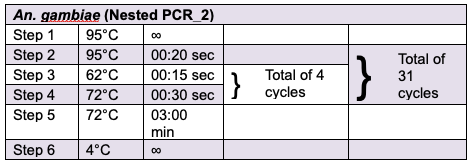  Temp cycles between 62/72C for 15.30s |
| PCR_1 | Primer number | 136 (68,66,2) | 80 |
| qPCR | Dilutions | 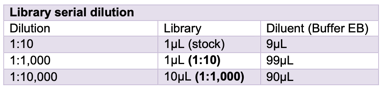  Dilutions 1:1k and 1:10k used | 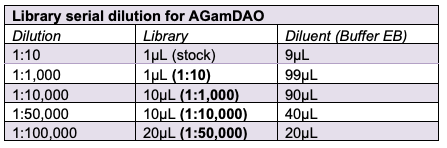  Additional dilutions required if PEP is used including 1:50k and 1:100k |
