## Supplementary Text 2 for "Targeted genomic surveillance of insecticide resistance in African malaria vectors"

### **Supplementary Text 2 - Ag-vampIR targets in detail**

The Ag-vampIR panel was designed as an extension of a previously published iPLEX MassARRAY panel (doi: 10.1038/s41598-019-49892-6), which comprised markers with known or suspected association with insecticide resistance and a set of markers to identify the different evolutionary origins of the Vgsc-995 pyrethroid target site mutations. The main additions to the initial panel are ancestry informative markers that can differentiate between *An. gambiae ss* and *An. coluzzii,* and markers that tag selective sweeps of potential interest for research into insecticide resistance. Genomic data from phase 1 of Ag1000G revealed the existence of several haplotypes undergoing recent positive selection in genes with known associations with resistance to insecticides, strongly suggestive that the haplotypes under selection confer increased resistance. We identified the haplotype under selection in each case, and included SNPs that were tagging the haplotypes in our panel. It will therefore be possible to track these haplotypes to monitor their frequency and geographical spread.

**Voltage-gated sodium channel (*Vgsc*)**

The voltage-gated sodium channel is the target site of pyrethroid insecticides and DDT. Mutations in this gene, particularly at codon 995, confer knockdown resistance (kdr) to these insecticides. We include both the L995F and L995S mutations, which have arisen independently multiple times across Africa. The L995F mutation is predominantly found in West Africa while L995S is more common in East Africa, though recent evidence suggests L995F is spreading eastward. We also include V402L and I1527T mutations that were recently discovered and are spreading in East Africa, along with the N1570Y mutation that enhances pyrethroid resistance when combined with L995F. Additionally, we target several non-synonymous mutations (V1853I, I1868T, P1874S/L, A1934V, A1746S, T791M and E1597G) that are strongly associated with L995F. These mutations may either enhance resistance or compensate for potential fitness costs of kdr. We also include 11 tagging SNPs across the *Vgsc* gene region that can differentiate between the different evolutionary origins of both L995F and L995S mutations, allowing the spread of specific kdr haplotypes to be tracked.

**Glutathione S-transferase epsilon cluster (Gste)**

The *Gste* gene cluster includes several genes involved in metabolic resistance to insecticides. We target the I114T mutation in Gste2 that has been shown to increase metabolism of DDT, as well as the L119V mutation which occurs at the same codon position as a mutation (L119F) conferring DDT resistance in *Anopheles funestus*. We also include tagging SNPs for multiple selective sweeps occurring in this gene cluster that may indicate novel resistance mechanisms.

**Cytochrome P450s**

**Cyp6p cluster**

The Cyp6p gene cluster contains several cytochrome P450 genes implicated in metabolic resistance. We target the I88T mutation in Cyp6p3, which is upregulated in resistant mosquitoes, and the I236M mutation in Cyp6p4 that tags a selective sweep and gene duplication in East African *An. gambiae*. Additional tagging SNPs are included to track multiple selective sweeps occurring across this cluster.

**Cyp9k1**

We include three tagging SNPs for selective sweeps occurring at the Cyp9k1 locus, which has been implicated in pyrethroid resistance through overexpression. These SNPs allow tracking of distinct swept haplotypes found in different populations.

**Other resistance-associated loci**

**Rdl**

The Rdl (resistance to dieldrin) or GABA gene contains mutations conferring resistance to cyclodienes. While these insecticides are no longer used for malaria control, the A296G and A296S mutations persist in natural populations. We include both mutations to track their frequencies and investigate potential cross-resistance to other compounds, which is likely as many insecticides target the GABA receptor.

**Ace1**

We target the G280S mutation in the acetylcholinesterase gene (Ace1) which confers resistance to organophosphates and carbamates. This mutation is often found as part of a heterogeneous duplication that combines resistant and susceptible alleles, potentially reducing fitness costs.

**Cyp4j5**

The L43F mutation in Cyp4j5 has been repeatedly associated with pyrethroid resistance in East Africa. In this region, the mutation is found at high frequency on the 2La inversion karyotype.

**Additional targets**

**Ancestry Informative Markers (AIMs)**

We include 35 ancestry informative markers that show fixed or nearly fixed differences between *An. gambiae* and *An. coluzzii*. These allow accurate species identification and detection of hybridization between these closely related species.

**TEP1**

We include two markers in the TEP1 gene which is involved in immune responses against *Plasmodium* parasites. While not directly related to insecticide resistance, these markers allow tracking of the TEP1*R1 allele associated with refractoriness to *Plasmodium falciparum*.

**34Mb selective sweep region**

We target four SNPs tagging a selective sweep at approximately 34Mb on chromosome arm 2L that is ongoing in *An. gambiae* populations from East Africa. While the functional significance of this sweep remains unknown, its rapid spread and association with the 2La inversion make it of potential interest for insecticide resistance.

**Doublesex**

We include two amplicons targeting the intron 4-exon 5 boundary of the doublesex gene, the target of a recently developed gene drive system for population suppression. These markers allow monitoring of any unintended spread of drive elements in wild populations.
