## Supplementary Text 3 for "Targeted genomic surveillance of insecticide resistance in African malaria vectors"

### **Supplementary Text 3 - AmpSeeker**

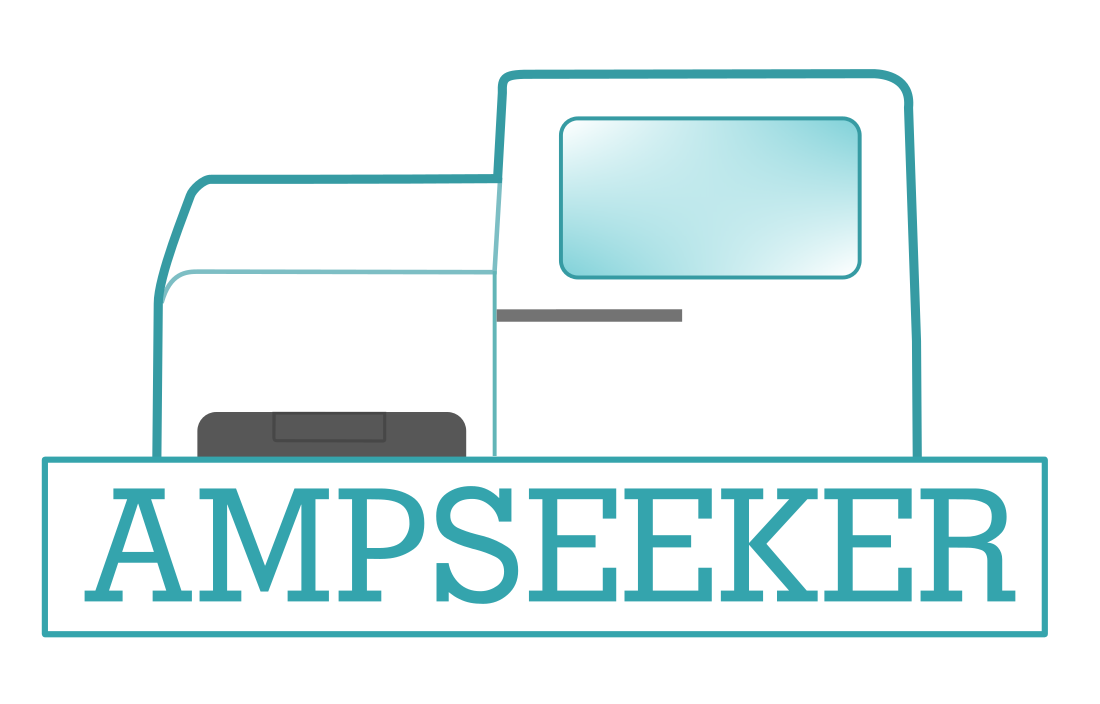

AmpSeeker is an open-source computational pipeline designed for the reproducible analysis of Illumina amplicon sequencing data. The pipeline provides end-to-end analysis capabilities, from raw sequencing data processing through to interactive visualization of results (Main text, Figure 1B). AmpSeeker implements best practices for amplicon sequence analysis while maintaining flexibility for different experimental designs and sequencing panels. While initially developed for the Ag-vampIR panel targeting insecticide resistance in *Anopheles gambiae sl,* AmpSeeker's architecture allows it to be applied to any Illumina amplicon sequencing data.

The workflow is built using Snakemake, a workflow management system that enables scalable and reproducible data analysis. Snakemake manages the execution of analysis steps through rules defined in separate files, with dependencies between steps automatically resolved through directed acyclic graphs. This ensures that analyses are performed in the correct order and that intermediate results are appropriately tracked.

The analytical components of AmpSeeker are implemented as Jupyter notebooks, which combine executable code, results, and documentation. These notebooks are executed programmatically using papermill, allowing parameters to be passed systematically while maintaining reproducibility. This architecture enables both automated execution through the pipeline and interactive exploration of results by users.

**AmpSeeker Folder Organization**

The AmpSeeker pipeline follows a standardized directory structure designed for clarity and reproducibility:

├── **config**

│	├── ag-vampir.bed

│	├── config.yaml

│	└── example-metadata.tsv

├── README.md

├── **resources**

│	├── ag-vampir

│	│	└── Kdr_marker_SNPs.csv

│	├── exampleSampleSheet.csv

│	├── multiqc.yaml

│	└── reference

└── **workflow**

├── ampseekertools.py

├── envs

│	├── AmpSeeker-cli.yaml

│	├── AmpSeeker-jupyterbook.yaml

│	├── AmpSeeker-python.yaml

│	├── AmpSeeker-qc.yaml

│	├── AmpSeeker-bcl2fastq.yaml

│	└── AmpSeeker-snpeff.yaml

├── notebooks

│	├── ag-vampir

│	│	├── kdr-analysis.ipynb

│	│	└── species-id.ipynb

│	├── allele-frequencies.ipynb

│	├── coverage.ipynb

│	├── genetic-diversity.ipynb

│	├── IGV-explore.ipynb

│	├── misc

│	│	├── primer-rebalancing.ipynb

│	│	└── reverse-complement.ipynb

│	├── population-structure.ipynb

│	├── process-notebooks.ipynb

│	├── process-toc.ipynb

│	├── read-quality.ipynb

│	├── reads-per-well.ipynb

│	├── run-information.ipynb

│	├── run-statistics.ipynb

│	├── sample-map.ipynb

│	├── sample-quality-control.ipynb

│	└── snp-dataframe.ipynb

├── **rules**

│	├── ag-vampir.smk

│	├── alignment-variantcalling.smk

│	├── analysis.smk

│	├── bcl-convert.smk

│	├── common.smk

│	├── jupyter-book.smk

│	├── qc-notebooks.smk

│	├── qc.smk

│	└── utilities.smk

└── **Snakefile**

10 directories, 43 files

The config directory contains the primary configuration file (config.yaml), metadata, and any panel-specific files such as bed files defining target regions. The resources directory holds reference files, including genome sequences, annotation files, and panel-specific resources. The workflow directory contains the core pipeline components:

- **envs**: Conda environment specifications for different analysis components

- **notebooks**: Jupyter notebooks implementing various analyses

- **rules**: Snakemake rule files defining the pipeline workflow

- **ampseekertools.py**: Utility functions used across the pipeline

Within the notebooks directory, analyses are organized by function, with panel-specific analyses (such as Ag-vampIR) in dedicated subdirectories. The rules directory similarly separates core functionality from panel-specific rules.

**Requirements and Dependencies**

AmpSeeker requires Python 3.8 or higher and manages software dependencies through conda. Currently, Conda, Snakemake and Pandas are required to begin the workflow. The pipeline is designed to run on Linux or MacOS systems and can be executed on both local machines, high-performance computing clusters and cloud-based systems through Snakemake's built-in support.

**Pipeline Modules and Outputs**

AmpSeeker consists of several core modules that process data sequentially:

- Read Processing and Quality Control processes raw sequencing data through BCL conversion, demultiplexing, and quality control with fastp. The module generates comprehensive quality metrics for both sequence and index reads.
- Alignment and Variant Calling performs sequence alignment using BWA-MEM, followed by variant calling with bcftools. The module includes coverage analysis with mosdepth and variant annotation using snpEff.
- Sample Quality Control implements filtering based on sequencing depth, missingness rates, and heterozygosity, generating detailed quality metrics and visualizations.
- Population Genetic Analysis performs various analyses including principal component analysis, neighbor-joining tree construction, and allele frequency analysis using scikit-allel.
- Each module generates standardized outputs organized in a consistent directory structure, with separate folders for raw data, processed data, quality control results, and analysis outputs.

**Local results web page (with Jupyter Book)**

AmpSeeker generates a comprehensive HTML report using Jupyter Book, which compiles all analysis notebooks into an interactive website. The report integrates quality control metrics, population genetic analyses, variant statistics, and interactive visualizations. This format allows users to explore results through an intuitive, familiar interface while maintaining access to the underlying analytical code.

The web page is organized hierarchically, with separate sections for different analyses and interactive visualizations implemented using Plotly, allowing the user to get a real feel for the data. The documentation includes methodological descriptions and guides to interpretation, making it accessible to both bioinformaticians and biologists.

**Extensibility**

AmpSeeker's modular architecture enables straightforward extension and customization. Users can add new analysis modules through additional Snakemake rules and Jupyter notebooks. The pipeline's use of standard file formats and interfaces facilitates integration with external tools and workflows. The separation of core pipeline functionality from panel-specific analyses allows AmpSeeker to be adapted for different experimental designs while maintaining its reproducible analysis framework. This extensibility ensures that AmpSeeker can evolve to analyse other species and meet new research needs while maintaining robust support for existing applications.
